# TRIM14 is a key regulator of the type I interferon response during *Mycobacterium tuberculosis* infection

**DOI:** 10.1101/828533

**Authors:** Caitlyn T. Hoffpauir, Samantha L. Bell, Kelsi O. West, Tao Jing, Sylvia Torres-Odio, Jeffery S. Cox, A. Phillip West, Pingwei Li, Kristin L. Patrick, Robert O. Watson

## Abstract

Tripartite motif-containing proteins (TRIMs) play a variety of recently described roles in innate immunity. While many TRIMs regulate type I interferon (IFN) expression following cytosolic nucleic acid sensing of viruses, their contribution to innate immune signaling and gene expression during bacterial infection remains largely unknown. Because *Mycobacterium tuberculosis* is a potent activator of cGAS-dependent cytosolic DNA sensing, we set out to investigate a role for TRIM proteins in regulating macrophage responses to *M. tuberculosis*. Here we demonstrate that TRIM14, a non-canonical TRIM that lacks an E3 ligase RING domain, is a critical negative regulator of the type I IFN response in macrophages. We show that TRIM14 physically interacts with both cGAS and TBK1 and that macrophages lacking TRIM14 dramatically hyperinduce interferon stimulated gene (ISG) expression following cytosolic nucleic acid transfection, IFN-β treatment, and *M. tuberculosis* infection. Consistent with a defect in resolution of the type I IFN response, *Trim14* knockout (KO) macrophages have more phospho-Ser754 STAT3 relative to phospho-727 and fail to upregulate the STAT3 target *Socs3* (Suppressor of Cytokine Signaling 3), which is required to turn off IFNAR signaling. These data support a model whereby TRIM14 acts as a scaffold between TBK1 and STAT3 to promote phosphorylation of STAT3 at Ser727 and enhance negative regulation of ISG expression. Remarkably, *Trim14* KO macrophages hyperinduce antimicrobials like *Inos2* and are significantly better than control cells at limiting *M. tuberculosis* replication. Collectively, these data reveal a previously unappreciated role for TRIM14 in resolving type I IFN responses and controlling *M. tuberculosis* infection.

## INTRODUCTION

*Mycobacterium tuberculosis*, arguably the world’s most successful pathogen, elicits a carefully orchestrated immune response that allows bacteria to survive and replicate in humans for decades. Infection of macrophages with *M. tuberculosis* sets off a number of pathogen sensing cascades, most notably those downstream of TLR2, which senses mycobacterial lipomannan (1, 2), and cGAS, which senses bacterial DNA in the host cytosol (3–5). cGAS-dependent DNA sensing during *M. tuberculosis* infection elicits two distinct and somewhat paradoxical responses: targeting of a population of bacilli for destruction in lysosomes via ubiquitin-mediated selective autophagy, and activation of a type I IFN gene expression program, which is inadequate at controlling bacterial pathogenesis *in vivo*. Because both selective autophagy and type I IFN have been repeatedly shown in animal and human studies to be hugely important in dictating *M. tuberculosis* infection outcomes (5–7), there is a critical need to elucidate the molecular mechanisms that drive their activation.

Many members of the TRIM family of proteins have emerged as important regulators of a variety of innate immune responses (6–8). Defined on the basis of their tripartite domain architecture, TRIMs generally encode a RING domain with E3 ligase activity, a B-box that is a zinc-binding domain with a RING-like fold (9), and a coil-coiled domain that mediates dimer/multimerization and protein-protein interactions (10). In addition to these domains, TRIMs encode a highly variable C-terminal domain. Since the initial discovery of TRIM5 as a potent HIV restriction factor (11), a variety of TRIMs have been shown to play critical roles in antiviral innate immunity through polyubiquitination of key molecules in DNA and RNA sensing cascades, including MDA5 by TRIM13, TRIM40, and TRIM65 (14–17), RIG-I by TRIM25 and TRIM40 (12, 13), and TBK1 by TRIM11 and TRIM23 (14). We are just beginning to appreciate the complex and dynamic network of regulatory factors, including TRIMs, that cells employ to up and downregulate innate immune signaling and gene expression (15).

Recent work has shown that cytosolic nucleic acid sensing pathways are also engaged during infection with a variety of intracellular bacterial pathogens, including *M. tuberculosis*, *Legionella pnuemophila*, *Listeria monocytogenes*, *Francisella novicida*, and *Chlamydia trachomatis* (16). Some of these pathogens, like *M. tuberculosis and C. trachomatis*, have been shown to activate cGAS via bacterial dsDNA (3, 17), while others like *L. monocytogenes* directly activate STING by secreting cyclic di-AMP (18). It is becoming increasingly clear that activation of cytosolic nucleic acid sensing pathways provides some benefit to intracellular bacterial pathogens, and thus the ability to engage with and manipulate regulatory molecules like TRIM proteins is likely a conserved bacterial adaptation. *Salmonella* Typhimurium has been shown to secrete SopA, an effector molecule which targets TRIM56 and TRIM65 to stimulate innate immune signaling through RIG-I and MDA5 (19, 20). Likewise, TRIM8 has been shown to regulate inflammatory gene expression downstream of TLR3 and TLR4 during *Salmonella* Typhimurium-induced septic shock (21). In addition, ablation of TRIM72 in alveolar macrophages enhances phagocytosis and clearance of *Psuedomonas aeruginosa* (22).

Realizing the huge potential for TRIM proteins in tipping the balance between pro- and anti-bacterial innate immune outcomes, we decided to study TRIMs during *M. tuberculosis* infection, specifically a non-canonical TRIM family member: TRIM14. Like most TRIMs, TRIM14 encodes a coiled-coil, a B-box, and a C-terminal PRY/SPRY domain, but curiously it lacks the E3 ligase RING domain, likely rendering it unable to catalyze ubiquitination of proteins. Consistent with it being a major player in antiviral innate immunity, TRIM14 has been shown to directly influence replication of several RNA viruses including influenza A via interaction with the viral NP protein (23), hepatitis B via interaction with HBx (24), and hepatitis C via interaction with NS5A (25). In the context of RNA sensing, TRIM14 has been shown to localize to mitochondria and interact with the antiviral signaling adapter MAVS (26). More recently, TRIM14 has been shown to promote cGAS stability by recruiting the deubiquitinase USP14 and preventing autophagosomal targeting of cGAS (27).

Here, we demonstrate that TRIM14 is a crucial negative regulator of *Ifnb* and ISG expression during macrophage infection with *M. tuberculosis*. TRIM14 is recruited to the *M. tuberculosis* phagosome and can directly interact with both cGAS and the DNA sensing kinase TBK1. Deletion of *Trim14* leads to dramatic hyperinduction of *Ifnb* and ISGs in response to *M. tuberculosis* and other cytosolic nucleic acid agonists. In *Trim14* KO macrophages we observe preferential phosphorylation of the transcription factor STAT3 at Ser754 and lack of association of STAT3 with the chromatin loci of target genes like Socs3, a crucial negative regulator of interferon α/β receptor (IFNAR) signaling. These data argue that TRIM14 acts as a negative regulator of cytosolic DNA sensing through bringing TBK1 and STAT3 together to promote phosphorylation of STAT3 at Ser727. Surprisingly, *Trim14* KO macrophages were remarkably efficient at limiting *M. tuberculosis* replication by virtue of overexpressing inducible nitric oxide synthase. Collectively, this work suggests that TRIM14 is a critical regulatory node of type I IFN induction and resolution in macrophages and highlights a previously unappreciated role for TRIM14 in anti-*M. tuberculosis* innate immunity.

## MATERIALS AND METHODS

### Cell Culture

RAW 264.7 macrophages, HEK293T, MEF, and LentiX cells were cultured at 37°C with 5% CO_2_. Cell culture medium was comprised of High glucose, sodium pyruvate, Dulbecco’s modified Eagle medium (Thermo Fisher) with 10% FBS (Sigma Aldrich) 0.5% HEPES (Thermo Fisher).

### Co-immunoprecipitations

1.8 × 10^6^ HEK293T cells in a 10cm plate were transfected with 1-10 μg of pDEST 3xFlag TRIM14, pDEST HA cGAS, pDEST HA STING, pDEST HA TBK1, pDEST HA IRF3, pDEST Flag STAT3, or pDEST HA TRIM14 using PolyJet In Vitro DNA Transfection Reagent. Cells were harvested in PBS+0.5M EDTA 24 hours post-transfection and pellets were lysed on ice with lysis buffer (50 mM Tris HCl pH 7.4, 150 mM NaCl, 1 mM EDTA, 0.075% NP-40) containing protease inhibitor (Pierce A32955). Strep-tactin superflow plus beads (Qiagen) were washed using buffer containing 5% 1M Tris at pH 7.4, 3% NaCl, and 0.2% 0.5M EDTA. 1000 μl of the cleared lysate was added to the beads and inverted for 2 hr at 4°C. Beads were then washed 4 times with wash buffer (50 mM Tris HCl pH 7.4 150 mM NaCl 0.5M EDTA, 0.05% NP-40) and eluted using 1x Biotin. Whole cell lysate inputs and elutions were boiled in 4x SDS loading buffer with 10% 2-mercapethanol. Proteins were run on SDS-PAGE gels (Bio-Rad) and then transferred to nitrocellulose membrane (GE Healthcare). Membranes were blocked in TBS (Odyssey Blocking buffer Li-COR) for 1 hour and incubated with primary antibody overnight at 4°C. LI-COR secondary was used (IR Dye CW 680 goat anti-rabbit, IR Dye CW 680 goat anti-rat 680, IR Dye CW800 goat anti-mouse (LI-COR)) and developed with the Odyssey Fc by LI-COR. Immunoprecipitation experiment were also performed as stated above but with Pierce Anti-HA agarose (Thermo 26181). Beads were eluted three times at room temperature for 15 min each using Influenza Hemagglutinin (HA) peptide (Sigma Aldrich I2149).

### Western Blot analysis

Protein samples were run on Any kD Mini-PROTEAN TGX precast protein gels (BioRad) and transferred to 0.45 μm nitrocellulose membranes (GE Healthcare). Membranes were incubated in the primary antibody of interest overnight and washed three times with TBS-Tween 20. Membranes were then incubated in secondary antibody for 1 hour and imaged using LI-COR Odyssey FC Imaging System. Primary antibodies used in this study: mouse monoclonal α-FLAG M2 antibody (Sigma-Aldrich, F3165), α-HA high affinity rat monoclonal antibody (Roche; 3F10), α-strep (Genscript A00626), α-phospho-Stat3 (Ser727) (Cell Signaling #9134), α-phospho-Stat3 (Ser754) (Cell Signaling #98543), α-Stat3 (124H6) Mouse mAb (Cell Signaling #9139), α-phospho-Stat1 (Tyr701) (58D6) Rabbit mAb #9167, α-TRIM14 G-15 (Santa Cruz sc79761), α-TRIM14 (Aviva ARP34737), and α-mouse monoclonal Beta-Actin (Abcam, #6276). Secondary antibodies used in this study: IR Dye CW 680 goat anti-rabbit, IR Dye CW 680 goat anti-rat 680, IR Dye CW800 goat anti-mouse (LI-COR), Alexfluor-488 anti-rabbit, Alexfluor-597 anti-rat, and Alexafluo-647 anti-mouse secondary antibodies for immunofluorescence (LI-COR).

### Construction of sgRNA/Cas9 LentiCRISPR and viral transduction

Guide RNAs targeting the first exon of Trim14 were designed using the Broad online tool (https://portals.broadinstitute.org/gpp/public/analysis-tools/sgrna-design). The top five hits were used to design five guide RNA constructs that were cloned into Lenti CRISPR vector (Puromycin) at the BsmB1 site. Constructs were sequenced and verified. Plasmids were then transfected into Lenti-X cells with PAX2 and VSVG packing plasmids. Virus was collected 24- and 48-hours post-transfection and stored at −80°C. RAW264.7 cells stably expressing Cas9 were then transduced with virus and selected using puromycin for 72 hours. Cells were then clonally selected using serial dilutions and clones were selected from wells calculated to contain a single cell. To verify the knockout, genomic DNA was collected from cells and the first exon of TRIM14 was amplified using PCR. This reaction was sent for sequencing to verify the mutation.

### Generation of shRNA-expressing stable cell lines

To generate knockdown RAW 264.7 macrophages, plasmids of scramble non-targeting shRNA constructs and TRIM14 shRNA constructs targeted towards the 3’ UTR of TRIM14 were transfected into Lenti-X cells with PAX2 and VSVG packing plasmids. Virus was collected 24 and 48-hours post-transfection and stored at −80°C. RAW264.7 cells were then transduced with virus and selected using hygromycin at 10mg/ml (Invitrogen) to select for cells containing the shRNA plasmid.

### Macrophage stimulation

RAW 264.7, CRISPR/Cas9 RAW 264.7, or shRNA RAW 264.7 macrophages were plated on 12- well tissue-culture treated plates at a density of 3 ×10^5^ cells/well and allowed to grow overnight. Cells were then transfected with 1 μg/ml ISD or 1 μg/ml poly(I:C) with lipofectamine or treated with 200units recombinant mouse IFNB (pbl assay science Cat#12400-1).

### *M. tuberculosis* Infection

Low passaged lab stocks of each Mtb strain (Erdman strain WT, Erdman *luxBCADE*, or Erdman m-Cherry) were thawed for each experiment to ensure virulence was preserved. *M. tuberculosis* was cultured in roller bottles at 37°C in Middlebrook 7H9 broth (BD Biosciences) supplemented with 10% OADC, 0.5% glycerol, and 0.1% Tween-80. All work with Mtb was performed under Biosafety Level 3 (BSL3) containment using procedures approved by the Texas A&M University Institutional Biosafety Committee. To prepare the inoculum, bacteria grown to log phase (OD 0.6-0.8) were spun at low speed (500g) to remove clumps and then pelleted and washed with PBS twice. Resuspended bacteria were briefly sonicated and spun at low speed once again to further remove clumps. The bacteria were diluted in DMEM + 10% horse serum and added to cells at an MOI of 10 for RNA and cytokine analysis and MOI of 1 for microscopy studies. Cells were spun with bacteria for 10 min at 1000 × g to synchronize infection, washed twice with PBS, and then incubated in fresh media. RAW 264.7 or CRISPR/Cas9 RAW 264.7 macrophages were plated on 12-well tissue-culture treated plates at a density of 3×10^5^ cells/well and allowed to grow overnight. Where applicable, RNA was harvested from infected cells using 0.5 ml Trizol reagent at each time point.

### *M. tuberculosis* survival/replication

RAW 264.7 or CRISPR/Cas9 RAW 264.7 macrophages were plated on 12-well tissue-culture treated plates at a density of 2.5 ×10^5^ cells per well. Luminescence was read for Mtb *luxBCADE* by lysing in 400ul 0.5% Triton-X and splitting into two wells of a 96 well white plate and using the luminescence feature of the INFINITE 200 PRO by TECAN at 0, 24, 48, and 72 hours post infection.

### RNA isolation and qPCR analysis

In order to analyze transcripts, cells were harvested in Trizol at the specified time points and RNA was isolated using Direct-zol RNA Miniprep kits (Zymo Research) with 1-hour DNase treatment. cDNA was synthesized with iScript cDNA Synthesis Kit (Bio-Rad). cDNA was diluted to 1:20 for each sample. A pool of cDNA from each treated or infected sample was used to make a 1:10 standard curve with each standard sample diluted 1:5 to produce a linear curve. RT-qPCR was performed using Power-Up SYBR Green Master Mix (Thermo Fisher) using a Quant Studio Flex 6 (Applied Biosystems). Samples were run in triplicate wells in a 96-well or 384 well plate. Averages of the raw values were normalized to average values for the same sample with the control gene, *beta-actin*. To analyze fold induction, the average of the treated sample was divided by the untreated control sample, which was set at 1.

### Immunofluorescence Microscopy

Glass coverslips were incubated in 100μl poly-lysine at 37°C for 30 minutes. MEF cells were plated at a density of 2×10^4^ on glass coverslips in 24-well plates and left to grow overnight. Cells were then transfected with 250ng of the desired plasmid(s) using PolyJet. The next day cells were treated with 1μg ISD as described above. At the designated time points, cells were washed with PBS (Thermo Fisher) and then fixed in 4% paraformaldehyde for 10 minutes. Fixed cells were washed three times in PBS and permeabilized by incubating them in PBS containing 5% non-fat milk and 0.05% saponin (Calbiochem). Coverslips were placed in primary antibody for 1 hour then washed 3x in PBS and placed in secondary antibody. These were washed twice in PBS and twice in deionized water, followed by mounting onto a glass slide using ProLong Diamond antifade mountant (Invitrogen). Images were acquired on a Nikon A1-Confocal Microscope. DAPI nuclear staining (Thermo Fisher).

### Co-localization experiments with *M. tuberculosis*

RAW 264.7 macrophages were plated on glass coverslips at a density of 3 ×10^5^ cells/well in 24-well plates. Cells were infected with m-Cherry *M. tuberculosis* at an MOI of 1 and fixed and stained as above at the designated time points. Colocalization of α-TRIM14 G-15 (Santa Cruz sc79761), with *M. tuberculosis* was visualized directly by fluorescence microscopy. A series of images were captured and analyzed by counting the number of bacteria that colocalized with the corresponding marker. At least one hundred events were analyzed per coverslip and each condition was performed with triplicate coverslips.

### Protein Expression and Purification

The cDNA encoding mouse TRIM14 (residues 247 to 440), mouse IRF-3 (residues 184-419) were cloned into a modified pET28(a) vector containing an N-terminal Avi-His6-SUMO tag. Sequences of the plasmids were confirmed by DNA sequencing. The BL21 (DE3) cells were co-transformed with the pET28(a) plasmids coding for the target proteins and the pBirAcm plasmid coding for BirA and induced with 0.4 mM IPTG in the presence of 5 μg ml−1 biotin and cultured at 16 °C overnight. The Biotin-labelled-Avi-His6-SUMO proteins were purified using a nickel-NTA column followed by gel-filtration chromatography using a HiLoad 16/60 Superdex 75 column (GE Healthcare). Mouse and human TBK1 (residues 1 to 657) were cloned into the pAcGHLTc baculovirus transfer vector. The plasmid was transfected together with Baculo-Gold bright linearized baculovirus DNA (BD Biosciences) into sf9 insect cells to generate recombinant baculovirus. The original recombinant viruses were amplified for at least two rounds before the large-scale protein expression. The insect cells at a density of 2.5 × 10^6^ cells/ml were infected by TBK1 recombinant baculovirus and cultured at 27°C and harvested 72 hours post infection. The cells were lysed in the buffer containing 150 mM NaCl, 0.2 M Tris-HCl, 1% NP-40, 1 mM PMSF at pH 8.0. The target protein in the supernatant was purified using nickel affinity chromatography followed by size-exclusion chromatography.

### SPR Binding Study

The binding studies between mouse TRIM14 and TBK1 were performed using a Biacore X100 SPR instrument (GE Healthcare). Biotin-labeled SUMO-fusion TRIM14 was coupled on the sensor chip SA (GE Healthcare). Dilution series of TBK1 or IRF-3 (1.25, 2.5, 5, 10, 20 μM) in 1× HBS-EP+ buffer (GE Healthcare) were injected over the sensor chip at a flow rate of 30 μL/min. The single-cycle kinetic/affinity protocol was used in all binding studies. All measurements were duplicated under the same conditions. The equilibrium K_d_ was determined by fitting the data to a steady-state 1:1 binding model using Biacore X100 Evaluation software version 2.0 (GE Healthcare).

### Chromatin Immunoprecipitation

Chromatin Immunoprecipitation (ChIP) was adapted from Abcam’s protocol. Briefly, one confluent 15 cm dish of CRISPR/Cas9 RAW 264.7 macrophages were crosslinked in formaldehyde to a final concentration of 0.75% and rotated for 10 minutes. Glycine was added to stop the cross linking by shaking for 5 minutes at a concentration of 125 mM. Cells were rinsed with PBS twice and then scraped into 5 mL PBS and centrifuged at 1,000g for 5 min at 4C. Cellular pellets were resuspended in ChIP lysis buffer (50 mM HEPES-KOH pH7.5, 140 mM NaCl, 1 mM EDTA pH8, 1% Triton X-100, 0.1% Sodium Deoxycholate, 0.1% SDS Protease Inhibitors) (750 μL per 1×10^7^ cells) and incubated for 10 min on ice. Cellular lysates were sonicated for 40 minutes (30sec ON, 30sec OFF) on high in a Bioruptor UCD-200 (Diagenode). After sonication, cellular debris was pelleted by centrifugation for 10 min, 4°C, 8,000 × g. Input samples were taken at this step and stored at −80°C until decrosslinking. Approximately 25 μg of DNA diluted to 1:10 with RIPA buffer was used for overnight immunoprecipitation. Each ChIP had one sample for the specific antibody and one sample for Protein G beads only which were pre-blocked for 1 hr with single stranded herring sperm DNA (75 ng/μL) and BSA (0.1 μg/μL). The respective primary antibody was added to all samples except the beads-only sample at a concentration of 5 ug and rotated at 4°C overnight. Beads were washed 3x in with a final wash in high salt (500mM NaCl). DNA was eluted with elution buffer and rotated for 15 min at 30C. Centrifuge for 1 min at 2,000 × g and transfer the supernatant into a fresh tube. Supernatant was incubated in NaCl, RNase A (10 mg/mL) and proteinase K (20 mg/mL) and incubated at 65°C for 1 h. DNA was purified using phenol:chloroform extraction. DNA levels were measure by RT-qPCR. Primers were designed by tiling each respective gene every 500 base pairs that were inputted into NCBI primer design.

### mRNA sequencing

RNA was sequenced from 4 biological replicates for each condition; Uninfected BMDMs, ESX-1-infected BMDMs, and M. tuberculosis-infected BMDMs. Raw reads were processed with expHTS (Street et al. 2015) to trim low-quality sequences and adapter contamination, and to remove PCR duplicates. Trimmed reads for each sample were aligned to the GRCm38 GENCODE primary genome assembly using STAR v.2.5.2b aligner (Dobin et al. 2013), and the GENCODE v.M10 annotation (gtf file). Each of the 4 replicates were merged into a single BAM file for further analysis. Prior to analysis, genes with expression less than 2 counts per million reads were filtered, leaving 11,808 genes. Differential gene expression was conducted using a single factor ANOVA model in the limma-voom Bioconductor pipeline. Log2 fold change values with a p-value <0.05 are represented in heatmaps where uninfected samples were the denominator and ESX-1 or *M.tuberculosis*-infected samples were the numerator in their respective datasets. Heatmaps were generated with GraphPad Prism Software.

### VSV infection

RAW 264.7 cells were seeded in 12-well plates at 8×10^5^ 16h before infection. Cells were infected with VSV-GFP virus at multiplicity of infection (MOI) of 1 in serum-free DMEM (HyClone SH30022.01). After 1h of incubation with media containing virus, supernatant was removed, and fresh DMEM plus 10% FBS was added to each well. At indicated times post infection, cells were harvested with Trizol and prepared for RNA isolation.

### Statistics

Statistical analysis of data was performed using GraphPad Prism software (GraphPad). Two-tailed unpaired Student’s t tests were used for statistical analyses, and unless otherwise noted, all results are representative of at least three independent biological experiments and are reported as the mean ± SD (n = 3 per group).

## RESULTS

### TRIM14 is a player in *M. tuberculosis* infection of macrophages

Having previously described a crucial role for the ESX-1 virulence-associated secretion system in eliciting cGAS-dependent cytosolic DNA sensing and type I IFN expression during *M. tuberculosis* infection, we first set out to better define gene expression differences in macrophages infected with a wild-type and a ΔESX-1 strain. Briefly, we infected bone marrow derived macrophages (BMDMs) with wild-type *M. tuberculosis* (Erdman strain) and the Tn5370::Rv3877/EccD1 mutant (ΔESX-1) (28), which lacks a functional ESX-1 secretion system and has previously been shown to be defective in eliciting cGAS-dependent responses (29, 30). We performed RNA-seq at an established key innate immune time point of 4h and an average log_2_ fold-change of 4 biological replicates is depicted (p < 0.05) (Fig. 1A). Consistent with previous microarray and RNA-seq data (30, 31), we observed dramatic induction of pro-inflammatory cytokines (*Il6*, *Il1b*) and antimicrobial molecules like *Inos2* in macrophages infected with both wild-type and ΔESX-1 *M. tuberculosis* (Fig. 1A), alongside downregulation of several protein-coding genes (*Epha2*, *Gpr34*, *Rtn4rl1*), and noncoding RNAs (Gm13391, Gm15564, Gm24270) (Fig. 1B). To identify genes/pathways whose induction requires ESX-1 secretion, we performed Ingenuity Pathway Analysis (Qiagen) and identified “Interferon signaling” and “Activation of IRF by Cytosolic PRRs” as the major pathways enriched for ESX-1 dependent genes (Fig. 1C and S1A-B). This analysis is in agreement with earlier data demonstrating a requirement for ESX-1 phagosome permeabilization for activation of type I IFN expression downstream of cGAS-dependent cytosolic DNA sensing (30). We next used RT-qPCR to measure expression of a number of important innate immune transcripts, both in BMDMs to validate our RNA-seq results and in RAW 264.7 murine macrophage-like cells (Fig. 1D and S1C), to justify our use of these genetically tractable cells moving forward. We observed almost identical expression of cytokines (*Ifnb*, *Il1b* (Fig. 1D), *Irf7*, Il6 (Fig. S1C)), and antimicrobial molecules (*Rsad2* (Fig. 1D), *Ifit*, *Isg15*, *Gbp1*, *Gbp5*, *Inos2* (Fig. S1C)) and in both cell types, maximal ISG induction was ESX-1-dependent while expression of NFκB genes was ESX-1-independent. In analyzing lists of ESX-1-dependent upregulated genes, we noticed that several belonged to the TRIM family, consistent with TRIMs being ISGs (Fig. 1E) (32). Because so little is known about how TRIM proteins regulate anti-bacterial immunity, we set out to better understand how TRIMs influence cGAS-dependent innate immune outcomes during *M. tuberculosis* infection.

**Figure 1:**
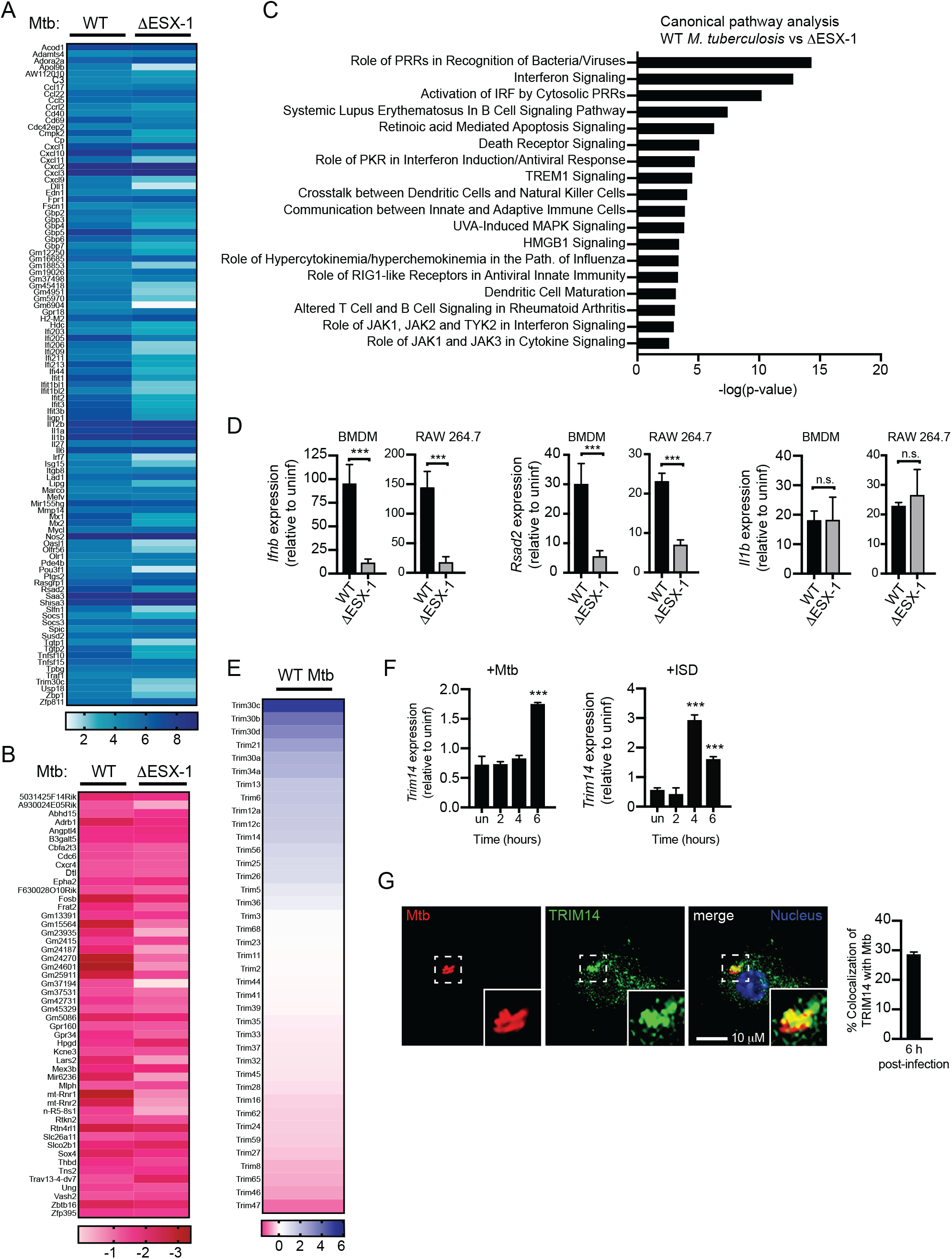
TRIMs are players in the innate immune response to *M. tuberculosis*. (A,B) Heatmap of significant (p < 0.05) gene expression differences (log_2_ fold-change over uninfected) in BMDMs infected with WT vs ΔESX-1 *M. tuberculosis* (Mtb) Genes upregulated are displayed in blue. Genes downregulated are displayed in red. (C) IPA software analysis of cellular pathways enriched for differentially expressed genes in BMDMs infected with WT vs ΔESX-1 *M. tuberculosis*. (D) RT-qPCR of transcripts in BMDMs & RAW 264.7 cells infected with WT *M. tuberculosis*. (E) Heatmap of significant (p < 0.05) gene expression differences (log_2_ fold-change) in TRIM family genes in BMDMS infected with WT vs ΔESX-1 *M. tuberculosis*. (F) RT-qPCR of fold-change in *Trim14* transcripts in BMDMs stimulated with ISD or infected with WT *M. tuberculosis*, n = 3 biological replicates. (G) RAW 264.7 cells infected with mCherry *M. tuberculosis* for 6 hours and immunostained for TRIM14. Statistical significance was determined using two-tailed students’ t test. *p < 0.05, **p < 0.01, ***p < 0.001.

Based on its recent characterization as a regulator of cGAS stability (27), we elected to investigate a role for TRIM14 during *M. tuberculosis* infection. RT-qPCR analysis confirmed that *Trim14* expression was upregulated after *M. tuberculosis* infection of RAW 264.7 cells (Fig. 1F). Transfection of RAW 264.7 cells with dsDNA (ISD) (33) recapitulated this effect (Fig. 1F), suggesting that *Trim14* upregulation during *M. tuberculosis* infection occurs downstream of cytosolic DNA sensing. To further implicate TRIM14 in *M. tuberculosis* infection of macrophages, we asked whether TRIM14 protein associated with the *M. tuberculosis* phagosome. Using immunofluorescence microscopy and an antibody against endogenous TRIM14, we detected TRIM14 at about 30% of *M. tuberculosis* phagosomes, reminiscent of the number of phagosomes we have previously shown to be positive for ubiquitin (Ub) and LC3, two markers of selective autophagy (Fig. 1G) (34). Together, these data begin to suggest that TRIM14 is a player in the macrophage response to *M. tuberculosis*.

### TRIM14 interacts with components of the DNA sensing pathway

Based on its recruitment to the *M. tuberculosis* phagosome, we hypothesized that TRIM14 may interact with one or more components of the cytosolic DNA sensing pathway (e.g. cGAS, TBK1) that we have previously observed to co-localize with *M. tuberculosis* (Fig. 2A and (3)). We transfected epitope-tagged versions of mouse TRIM14 (3xFLAG-TRIM14) and major components of the DNA sensing pathway (mouse HA-cGAS, HA-STING, and HA-TBK1) into murine embryonic fibroblasts (MEFs). Following 24 hours of expression, cells were fixed, and co-immunostained. Consistent with a previous report (24), we observed that TRIM14 co-localized with cGAS (Fig. 2B), while no co-localization between 3xFLAG-TRIM14 and HA-STING was seen. Intriguingly, we also detected substantial overlap between 3xFLAG-TRIM14 and HA-TBK1, suggesting that TRIM14 may interact with more than one component of the cytosolic DNA sensing pathway.

**Figure 2:**
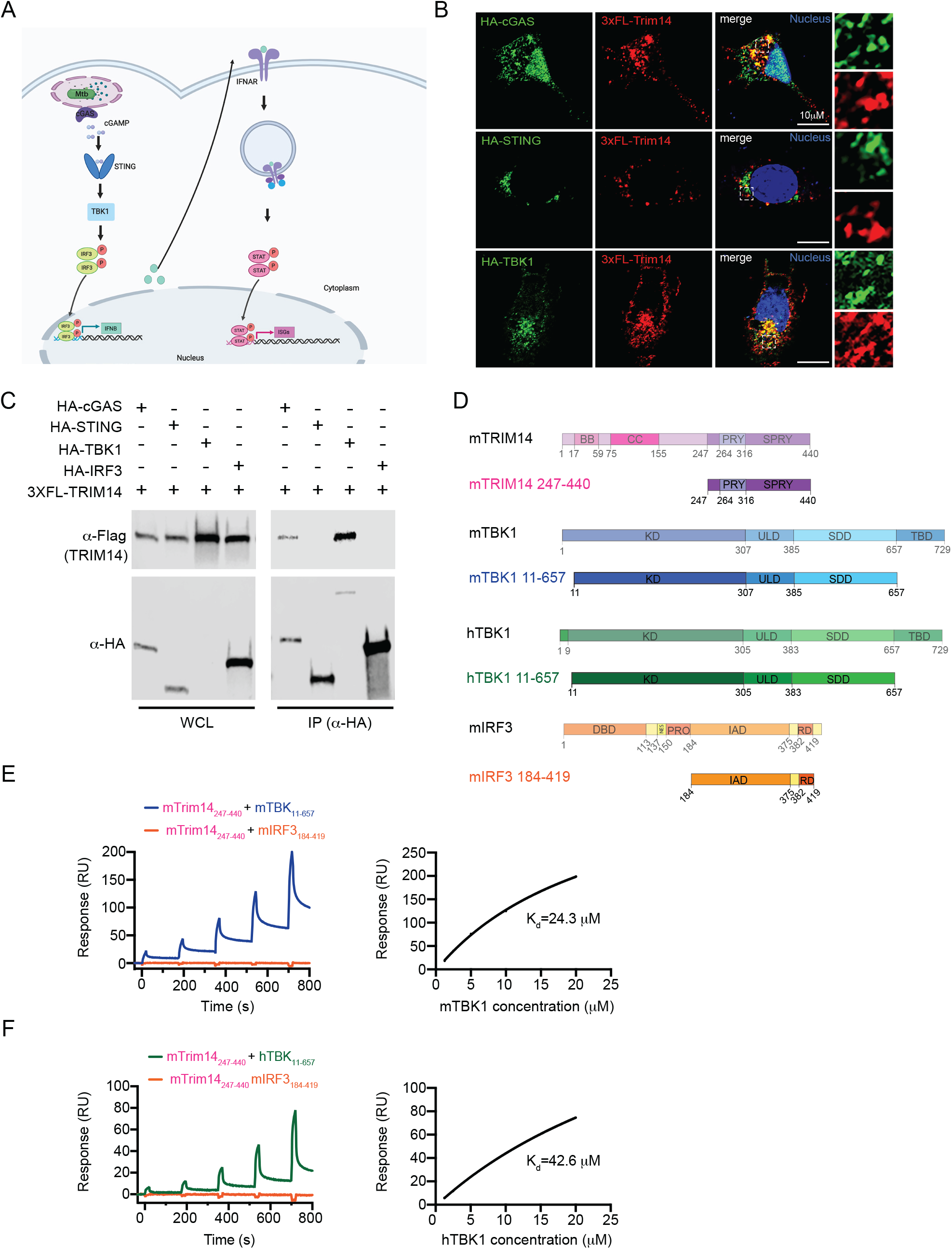
TRIM14 interacts with both cGAS and TBK1 in the DNA sensing pathway. (A) Model of DNA sensing pathway during *M. tuberculosis* infection (B) Immunofluorescence microscopy of MEFs expressing 3xFLAG-TRIM14 with HA-cGAS, HA-STING, or HA-TBK1 co-stained with α-Flag antibodies (C) Western blot analysis of co-immunoprecipitation of 3xFLAG-TRIM14 co-expressed with HA-cGAS, HA-STING, HA-TBK1, or HA-IRF3 in HEK 293T cells. Blot is representative of >3 independent biological replicates. (D) Diagram of mTRIM14, mTBK1, hTBK1, mIRF3 gene domains and truncations used in surface plasmon resonance (SPR) studies. (E) Equilibrium binding study of mTRIM14 and mTBK1 by SPR. mIRF3 was used as a negative control. Dissociation constant (K_d_= 24.3μM) was derived by fitting of the equilibrium binding data to a one-site binding model. (F) As in (E) but with mTRIM14 and hTBK1. Dissociatio constant (K_d_= 42.6μM) was derived by fitting of the equilibrium binding data to a one-site binding model.

To further characterize this previously unappreciated association between TRIM14 and TBK1, we co-expressed mouse 3xFLAG-TRIM14 with mouse HA-cGAS, HA-STING, HA-TBK1, and HA-IRF3 in HEK 293T cells, immunopurified each of the DNA sensing pathway components, and probed for interaction with TRIM14 by western blot. Consistent with our immunofluorescence microscopy data, we found that TRIM14 co-immunoprecipitated with both cGAS and TBK1 but not STING or IRF3 (Fig. 2C).

Next, to determine whether these biochemical associations were direct interaction, we performed surface plasmon resonance (SPR) experiments. Briefly, truncated versions of mouse TRIM14 (residues 247-440), human cGAS (residues 157-522), and mouse and human TBK1 (residues 11-657) were expressed using a baculovirus system. A portion of mouse IRF3 (residues 184-419) served as the negative control (Fig. 2D). Each of these protein truncations had previously been shown to be stably expressed at high levels and remain soluble when generated in insect cells ((35–37) and Fig. S2A). Equilibrium binding studies measured a binding affinity of 24.3 μM for binding between mTRIM14 and mTBK1 (Fig. 2E) and a slightly lower affinity of 42.6 μM for mouse TRIM14 and a portion of human TBK1 (Fig. 2F). Mouse TRIM14 and full-length hTBK1 also showed direct binding (K_d_ = 11 μM (Fig. S2D)), as did human cGAS and mouse TRIM14 (K_d_ = 25.8 μM (Fig. S2B-C)). No binding was measured between mTRIM14 and mIRF3 in any of the experiments (Fig. 2E-F and S2C-D). Combined, these *in vivo* and *in vitro* biochemical data argue strongly for a direct interaction between TRIM14 and TBK1.

### Loss of TRIM14 leads to type I IFN and ISG hyperinduction during *M. tuberculosis* infection

To investigate the contribution of TRIM14 to cytosolic DNA sensing outcomes during *M. tuberculosis* infection, we first tested how knockdown of *Trim14* affects *Ifnb* gene expression. *Trim14* knockdown (KD) macrophages were generated by transducing RAW 264.7 cells with lentiviral shRNA constructs designed to target the 3’UTR of *Trim14* or a control scramble shRNA (SCR). RT-qPCR analysis confirmed ~50% and 70% knockdown of *Trim14* using two different shRNA constructs (KD1 and KD2 respectively) (Fig. 3A). *Trim14* KD and control RAW 264.7 cells were either infected with *M. tuberculosis* or transfected with ISD to directly engage cGAS and *Ifnb* transcripts were measured after 4 hours. In both experiments, we observed lower levels of *Ifnb* transcript induction in *Trim14* KD cell lines compared to the SCR control (Fig. 3B and 3C), supporting a role for TRIM14 in the DNA sensing pathway. Since residual levels of TRIM14 protein in knockdown cell lines could potentially complicate interpretation of phenotypes, we decided to generate *Trim14* knockouts (KO) using CRISPR-Cas9. Briefly, *Trim14*-specific guide RNAs (gRNAs) were designed to target *Trim14* exon 1 and a GFP-specific gRNA was designed as a negative control. Two clones with distinct frameshift mutations that each introduced a stop codon early in exon 1 were identified and chosen for subsequent experimentation (Fig. 3D). Knockout of *Trim14* was confirmed by western blot using an antibody against the endogenous protein and by anti-TRIM14 immunofluorescence of control and *Trim14* KO cells (Fig. 3E).

**Figure 3:**
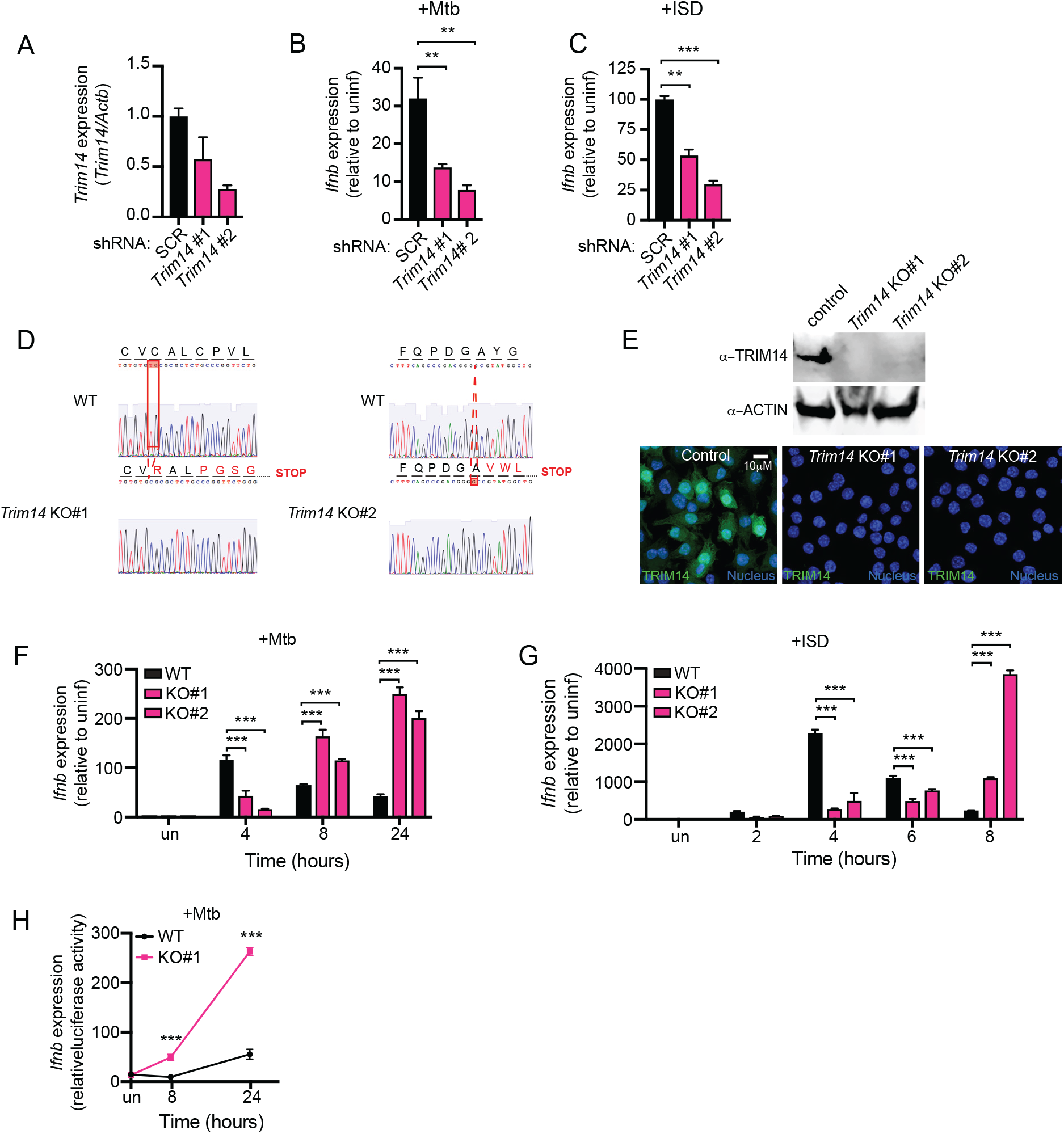
Loss of Trim14 leads to hyperinduction of *Ifnb* in response to *M. tuberculosis* and cytosolic DNA. (A) RT-qPCR of *Trim14* transcript in RAW 264.7 macrophages stably expressing shRNA to either SCR control or *Trim14*. (B) RT-qPCR of *Ifnb* transcript in RAW 264.7 macrophages stably expressing shRNA to either SCR control or *Trim14* infected with *M. tuberculosis* at 4h post infection. (C) RT-qPCR of *Ifnb* transcript in RAW 264.7 macrophages stably expressing shRNA to either SCR control or TRIM14 transfected with 1μg ISD at 4h post infection. (D) Sequencing chromatogram depicting mutations in *Trim14* gRNA CRISPR/Cas9 RAW 264.7 macrophages compared to GFP gRNA control (WT). (E) Western blot analysis and immunofluorescence microscopy of *Trim14* in WT vs *Trim14* KO RAW 264.7 macrophages using an anti-TRIM14 antibody. (F) RT-qPCR of *Ifnb* transcripts in WT and *Trim14* KO RAW 264.7 macro-phages infected with *M. tuberculosis* at specified times after infection. (G) RT-qPCR of *Ifnb* transcripts in WT and *Trim14* KO RAW 264.7 microphages treated with ISD at specified times after treatment. (H) ISRE reporter cells expressing luciferase with relative light units measured as a readout for IFN-β protein. All RT-qPCRs represent n = 3 biological replicates. Statistical significance was determined using two-tailed students’ t test. *p < 0.05, **p < 0.01, ***p < 0.001.

In order to test how genetic ablation of *Trim14* affects cytosolic DNA sensing, *Trim14* KO and control macrophages were infected with wild type *M. tuberculosis* and RNA was collected over a 24h time-course of infection. Surprisingly, while we again measured lower *Ifnb* expression at 4h post-infection, we observed a dramatic hyperinduction of *Ifnb* in the absence of TRIM14 at later infection time points (Fig. 3F). To determine the contribution of cytosolic DNA sensing to this phenotype, we transfected *Trim14* KO and control RAW 264.7 cells with ISD and again found significantly higher induction of *Ifnb* in the absence of TRIM14 at 8 hours and 24 hours post-transfection (Fig. 3G). To verify that the transcript changes we observed translated to differences in protein levels, we used Interferon Stimulated Response Element (ISRE) luciferase reporter cells to analyze IFN-β protein secretion in supernatants from cells 8- and 24-hours post-*M. tuberculosis* infection. Using relative light units as a proxy for IFN-β, we again observed higher production of IFN-β in the absence of TRIM14 (Fig. 3H). These data suggest that the major phenotype associated with *Trim14* ablation in RAW 264.7 macrophages is loss of negative regulation following cytosolic DNA sensing.

Having observed higher *Ifnb* transcript and protein levels in *Trim14* KO RAW 264.7 cells, we predicted that these cells would also hyperinduce ISGs following treatment with innate immune agonists that stimulate IRF3 signaling downstream of cGAS or STAT signaling downstream of IFNAR. RT-qPCR analysis of RNA recovered over a time-course of either *M. tuberculosis* infection or ISD transfection showed hyper induction of *Ifit1*, *Isg15*, and *Irf7* (Fig. 4A and B). Likewise, high levels of these same ISGs were observed when cells were transfected with 1μg poly (I:C), a potent agonist of RNA sensing via RIG-I (Fig. 4C) or treated with recombinant IFN-β directly (Fig. 4D). Importantly, non-ISGs like *Il1b*, *Tnf*, and *Il6*, despite being dramatically induced during *M. tuberculosis* infection were completely unaffected by loss of TRIM14 (Fig. S3A). Collectively, these results demonstrate that TRIM14 is needed for induction of *Ifnb*/ISGs immediately following cGAS sensing as well as for subsequent resolution of the response, arguing for a model whereby TRIM14 regulates cytosolic DNA sensing at two distinct nodes in the pathway.

**Figure 4:**
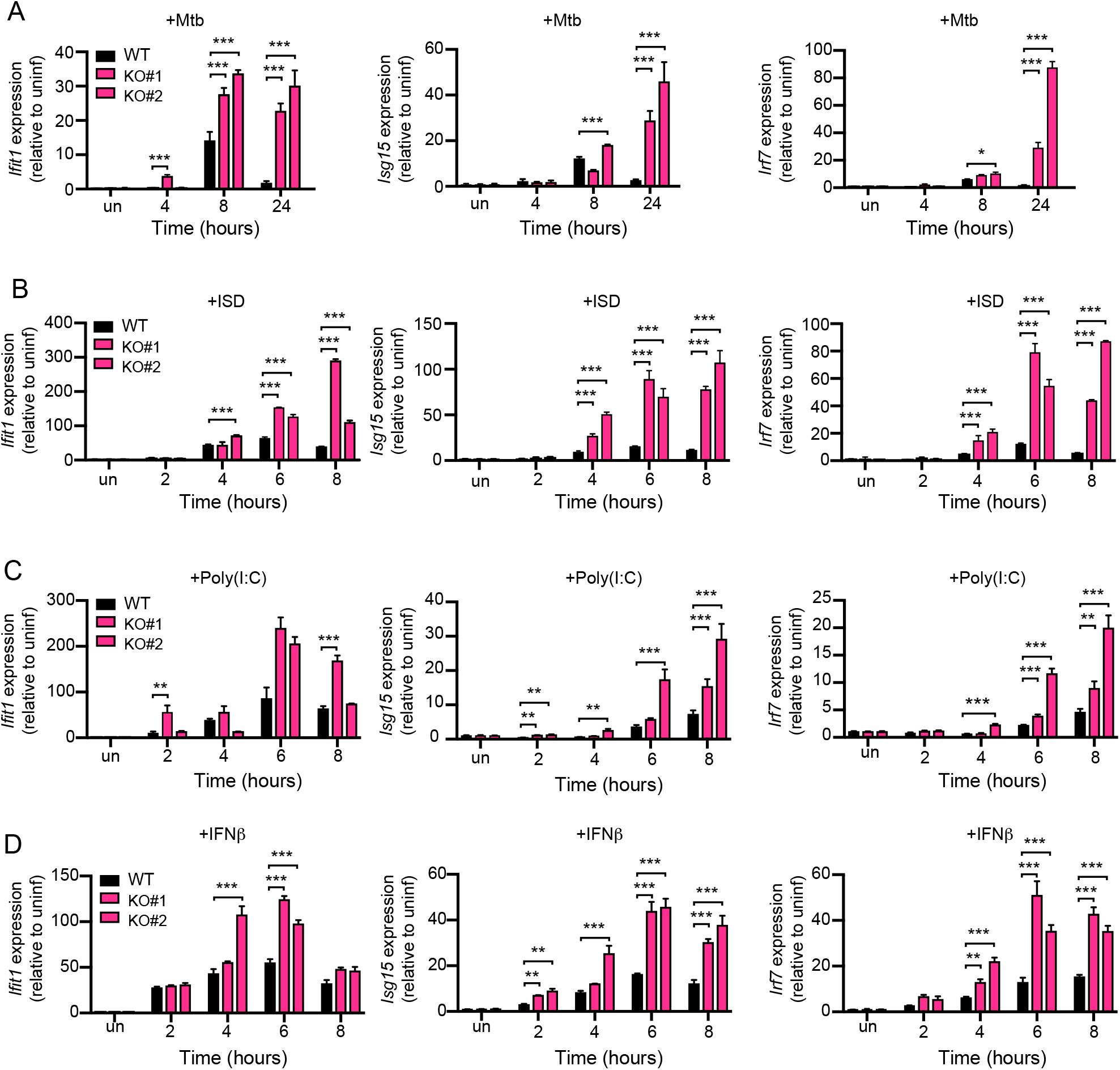
Loss of TRIM14 leads to hyperinduction of ISGs in response to multiple innate immune agonists. (A) RT-qPCR of *Ifit*, *isg15*, *Irf7* transcripts in WT and *Trim14* KO RAW 264.7 macrophages infected with *M. tuberculosis* at specified times after infection. (B) RT-qPCR of *Ifit*, *Isg15*, *Irf7* transcripts in WT and *Trim14* KO RAW 264.7 macrophages transfected with 1μg ISD. (C) As in (B) but with poly(I:C) transfection. (D) As in (B) but with IFNβ treatment (200 IU). All RT-qPCRs represent n = 3 biological replicates. Statistical significance was determined using two-tailed students’ t test. *p < 0.05, **p < 0.01, ***p < 0.001.

### TRIM14 regulates STAT3 activation through TBK1

Since we detected *in vivo* and *in vitro* interactions between TRIM14 and TBK1, we hypothesized that TRIM14-dependent misregulation of TBK1 activity could drive hyperinduction of *Ifnb* and ISGs. TBK1 is a prolific innate immune serine/threonine kinase with many known targets (38, 39). One such target, STAT3 (Signal transducer and activator of transcription 3) has been repeatedly implicated in negatively regulating type I IFN responses (40, 41). Therefore, we set out to determine whether the presence of TRIM14 and its interaction with TBK1 was required to control STAT3 activity in macrophages.

Previous studies have demonstrated that TBK1 can directly phosphorylate STAT3 at Ser727 and Ser754 upon cytosolic DNA sensing (45) (Fig. 5A). Phosphorylation of STAT3 at Ser754 inhibits STAT3’s ability to interact with target genes, while phosphorylation of STAT3 at Ser727 promotes STAT3 activity and transcription of STAT3 targets (42). To determine whether TRIM14 influences STAT3 phosphorylation, we transfected control and *Trim14* KO cells with ISD to activate TBK1 and analyzed STAT3 phosphorylation at Ser727 and Ser754 by immunoblot over a time-course. We observed substantially more phospho-Ser754 STAT3 and significantly less phospho-Ser727 STAT3 in the absence of TRIM14 (Fig. 5B; replicate blot in Fig. S4B), while loss of TRIM14 had no effect on JAK tyrosine kinase phosphorylation of STAT1 at Y701. These data suggest a role for TRIM14 in influencing TBK1’s preference to phosphorylate particular serine residues in the transactivation domain of STAT3. To further implicate TRIM14 in mediating STAT3 activation by TBK1, we performed cellular fractionation experiments and measured the amount of STAT3 in the nucleus following ISD transfection in *Trim14* KO and control cells by immunoblot. Consistent with reduced activation of STAT3, we observed significantly less STAT3 accumulation in the nuclei of *Trim14* KO cells (Fig. 5C). We next predicted that TRIM14 can control TBK1’s ability to phosphorylate STAT3 by interacting with both factors and bringing them together in a conformation that promotes phosphorylation at STAT3 S727 while inhibiting phosphorylation at S754. Previous studies have demonstrated that TBK1 and STAT3 can co-immunoprecipitate (42). To “close the loop”, we tested whether STAT3 and TRIM14 can interact when co-expressed in HEK293T cells and indeed, detected association between STAT3 and TRIM14 (Fig. 5D)., We propose that TRIM14, through interactions with STAT3 and TBK1, influences TBK1’s ability to phosphorylate STAT3 at particular serine residues.

**Figure 5:**
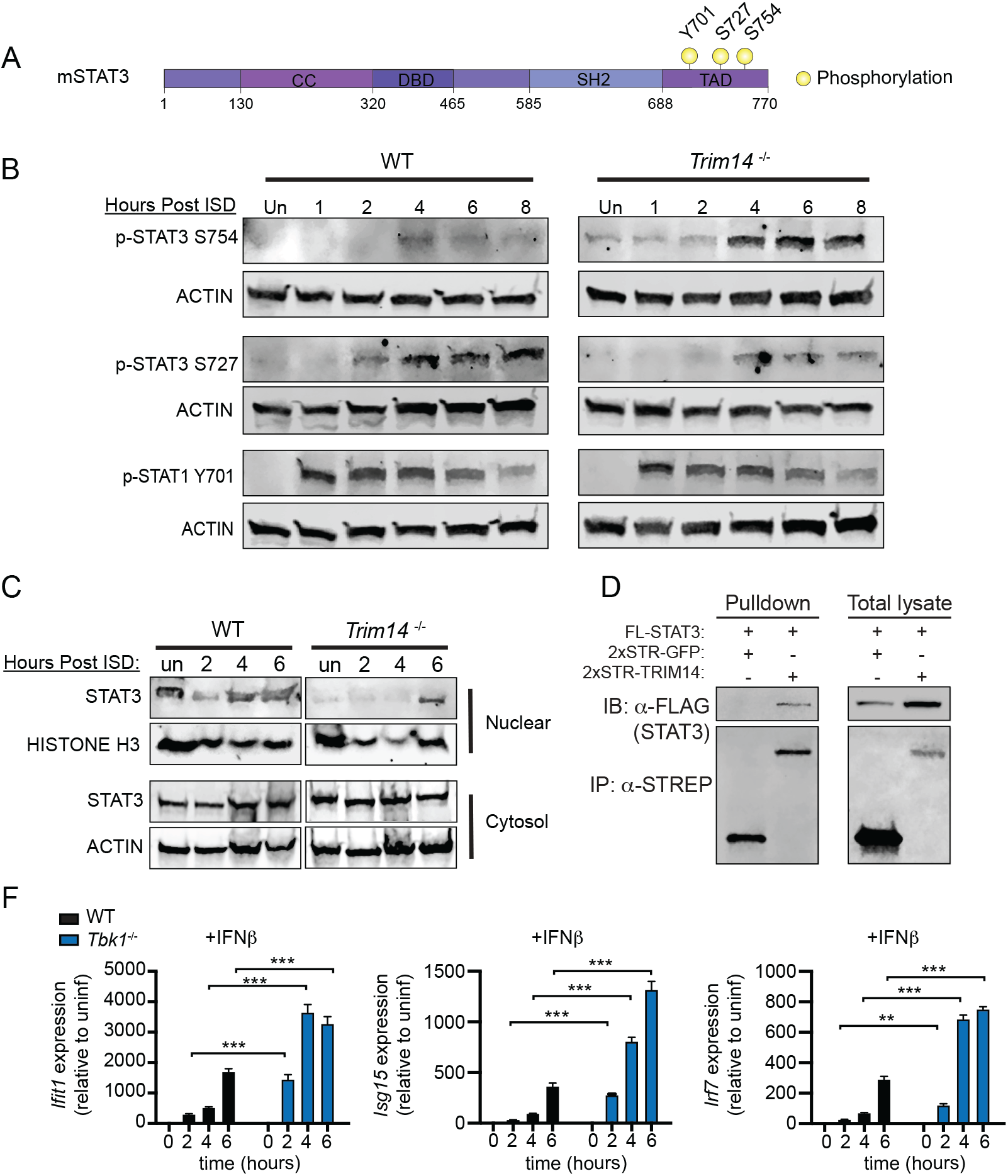
Loss of TRIM14 promotes inhibitory phosphorylation of STAT3 at Ser754. (A) Cartoon depiction of phosphorylation sites of STAT3. (B) Immunoblot of phospho-STAT3 Ser754, phospho-STAT3 Ser727, and phospho-STAT1 Y701 in WT and *Trim14* KO RAW 264.7 macrophages at 1, 2, 4, 6, 8h post-ISD transfection. ACTIN is shown as a loading control. (C) Cellular fractionation showing nuclear STAT3 in WT and *Trim14* KO RAW 264.7 macrophages treated with ISD at specified times after treatment. Histone 3 shown as loading/nuclear control (D) Co-immunoprecipitation and western blot analysis of HEK 293T cells co-transfected with FLAG-STAT3 and HA-TRIM14. (E) RT-qPCR of *Ifit*, *Isg15*, *Irf7* transcripts in WT and *Tbk1* KO BMDMs treated with IFN-β. Western blots are representative of >3 independent biological replicates. Statistical significance was determined using two-tailed students’ t test. *p < 0.05, **p < 0.01, ***p <0.001.

We reasoned that since loss of TRIM14 caused hyperinduction of ISGs via a TBK1/STAT3 dependent mechanism, then loss of either TBK1 or STAT3 would phenocopy loss of TRIM14. Indeed, previous studies have shown that *Stat3* KO MEFs and BMDMS hyperinduce ISGs following viral infection (40). To test whether loss of TBK1 could also lead to ISG hyperinduction, we harvested BMDMs from *Tbk1*^−/−^/*Tnfr*^−/−^ mice (43, 44) and treated them with recombinant IFN-β to directly engage with IFNAR and bypass the need for TBK1 to phosphorylate IRF3 and promote *Ifnb* expression (45, 46). Remarkably, we measured dramatic hyperinduction of ISGs in *Tbk1*^−/−^/*Tnfr*^−/−^ BMDMs over a six-hour time course of IFN-β treatment (Fig. 5F). This result argues that TBK1 plays a crucial, yet mostly unappreciated, role in diminishing the type I IFN response downstream of IFNAR signaling and supports a model whereby TRIM14 downregulates type I IFN gene expression via TBK1-dependent phosphorylation of STAT3.

### TRIM14 is required for STAT3-dependent transcription of *Socs3*, a negative regulator of the type I IFN response

Because an uncontrolled type I IFN response is deleterious to the host, cells have evolved multiple mechanisms to dampen type I IFN gene expression following pathogen sensing or IFNAR activation. The hyperinduction of *Ifnb* and ISGs we measure in *Trim14* KO macrophages is consistent with a loss of negative regulation; therefore, we hypothesized that expression of one or more negative regulators would be lower in *Trim14* KO cells. Inhibition of JAK1-STAT signaling is a well-characterized mechanism through which type I IFN signaling is downregulated (47). SOCS (Suppressor of cytokine signaling) family proteins are ISGs that dampen type I IFN responses by interfering with JAK1 kinase activity and limiting STAT signaling (41). USP18 (Ubiquitin Specific Peptidase 18) has similarly been shown to inhibit type I IFN expression by blocking interaction between JAK1 and the IFNAR2 subunit (48). To test whether *Trim14* KO cells were defective in expressing negative regulators of the type I IFN response, we measured *Socs3*, *Socs1*, and *Usp18* transcripts in control and *Trim14* KO RAW 264.7 cells infected with *M. tuberculosis* or transfected with ISD. Similar to all ISGs we examined in these studies, *Socs1* and *Usp18* were hyperinduced in *Trim14* KO macrophages. However, we observed a specific defect in *Socs3* induction, suggesting that one or more *Socs3* transcription factors were impacted by loss of TRIM14 (Figure 6A and 6B).

**Figure 6:**
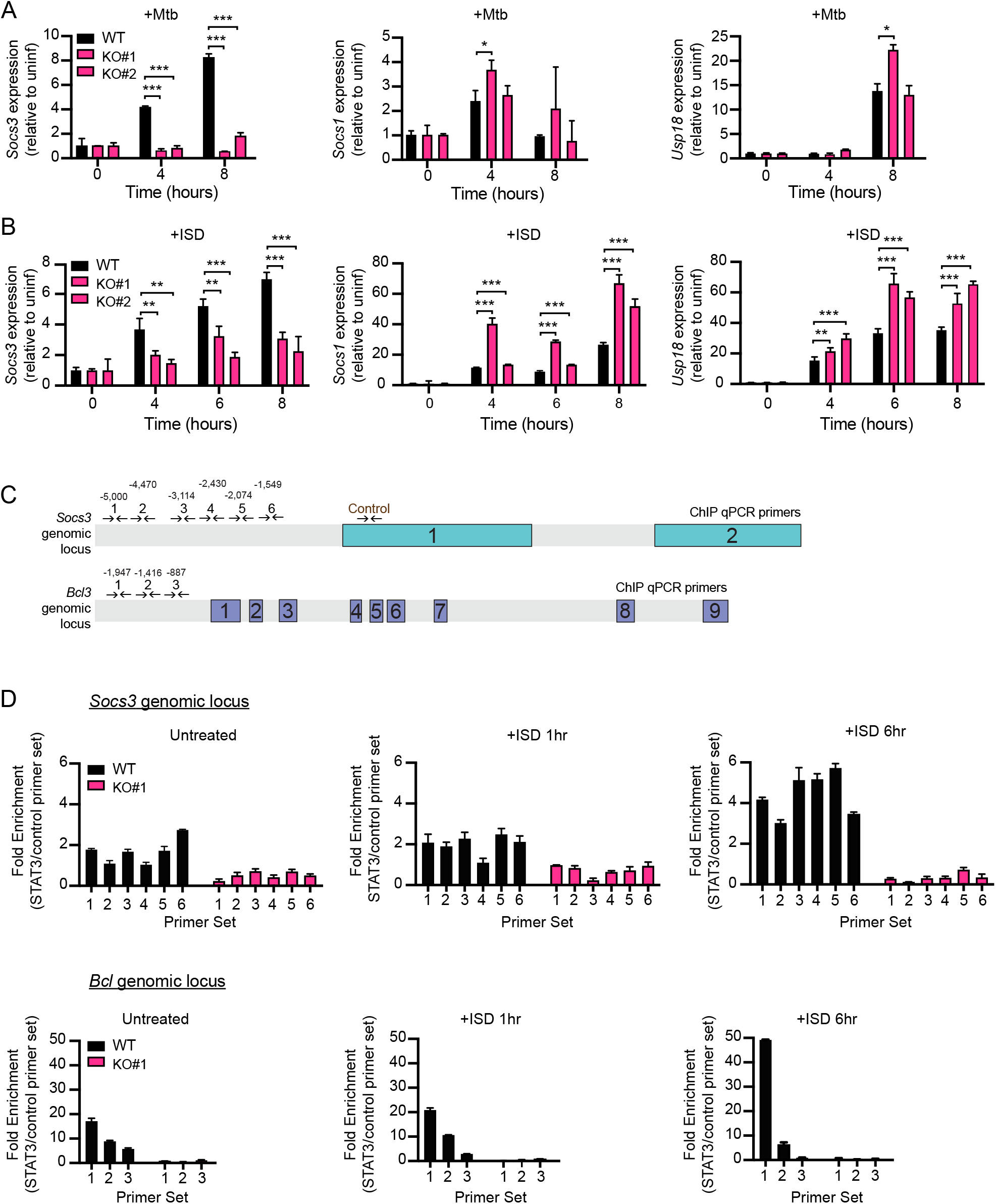
*Trim14* KO macrophages fail to induce expression of the negative regulator of the type I IFN response, Socs3. (A) RT-qPCR of *Socs3*, *Socs1*, and *Usp18* transcripts in WT and *Trim14* KO RAW 264.7 macrophages infected with *M. tuberculosis* at specified times after treatment. (B) RT-qPCR of *Socs3*, *Socs1*, and *Usp18* transcripts in WT and *Trim14* KO RAW 264.7 macrophages treated with ISD at specified times after treatment. (C) qPCR primers designed to tile *Socs3* and *Bcl3* genes for chromatin immunoprecipitation experiments (D) ChIP-qPCR of 3xFLAG-STAT3 associated genomic DNA from the *Socs3* and *Bcl3* loci in WT and *Trim14* KO RAW 264.7 macrophages transfected with 1μg ISD. Statistical significance was determined using two-tailed students’ t-test. *p < 0.05, **p < 0.01, ***p < 0.001.

Previous reports have shown that *Socs3* is a major target gene of STAT3 (49, 50). Having measured increased phosphorylation at the “inhibitory” Ser754 residue of STAT3 in *Trim14* KO macrophages, we hypothesized that lack of *Socs3* induction could be due to the inability of STAT3 to bind at the *Socs3* promoter. To test this, we transfected control and *Trim14* KO cells with ISD and performed chromatin immunoprecipitation (ChIP) at 0h, 1h, and 6h following ISD transfection using an antibody directed against total STAT3 protein. Consistent with low *Socs3* transcription, we detected significantly less recruitment of STAT3 to the *Socs3* genomic locus at both 1h and 6h after ISD transfection (Fig. 6D). We also detected lower STAT3 recruitment to other non-ISG target genes, including *Bcl3* (Fig. 6D) and *Cxcl9* (Fig. S3C), suggesting that loss of TRIM14 broadly impacts STAT3’s ability to translocate to the nucleus and/or associate with DNA. From these data, we conclude that defective nuclear translocation of STAT3 and subsequent lack of *Socs3* transcriptional activation result in ISG hyperinduction in the absence of TRIM14.

### Loss of TRIM14 impacts the ability of macrophages to control infection

Having demonstrated an important role for TRIM14 in regulating the type I IFN response, we set out to investigate how loss of TRIM14 impacts cell-autonomous innate immune responses to viral and intracellular bacterial infection. To test how loss of TRIM14 impacts survival and replication of *M. tuberculosis*, we infected control and *Trim14* KO macrophages with *M. tuberculosis* expressing the *luxBCADE* operon from *Vibrio harveyi* (51) and quantified bacterial replication as a measure of luminescence over a 72 hour time course (51, 52). Remarkably, we observed a dramatic inhibition of *M. tuberculosis* replication in *Trim14* KO macrophages (Figure 7A). Importantly, this lack of *M. tuberculosis* replication did not correspond to loss of cells, as infected monolayers remained completely intact at the 72h time point (Fig. S4A). We also observed a significant inhibition of *M. tuberculosis* replication in *Trim14* KO RAW264.7 cells by enumerating colony forming units (CFUs) (Fig. 7B). To begin to identify the molecular mechanisms driving *M. tuberculosis* restriction in *Trim14* KO macrophages, we measured expression of several genes whose proteins have been purported to have direct bactericidal activity against *M. tuberculosis* (53, 54, 55). RT-qPCR revealed hyperinduction of inducible nitric oxide (*Inos2*) and guanylate binding proteins 1 and 5 (*Gbp1* and *Gbp5*) in *M. tuberculosis*-infected *Trim14* KO macrophages (Fig. 7C). We predict that the overabundance of one or more of these factors contributes to enhanced control of *M. tuberculosis* replication in the absence of TRIM14.

**Figure 7:**
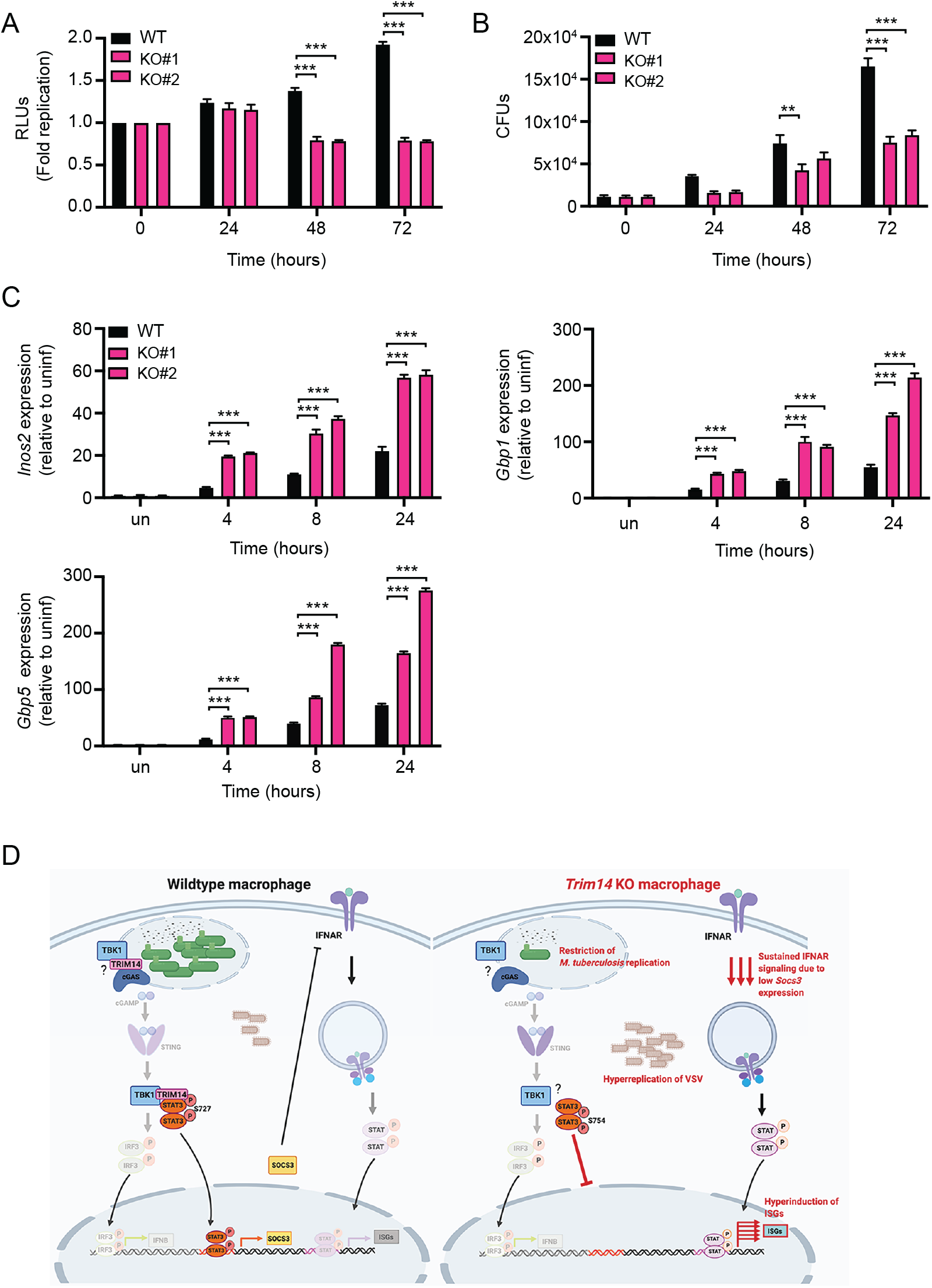
*Trim14* KO macrophages restrict *M. tuberculosis* replication but fail to control VSV replication. (A) Fold replication of *luxBCADE* in WT and *Trim14* KO RAW 264.7 macrophages. (B) CFUs in WT and *Trim14* KO RAW 264.7 macrophages infected with at 0h, 24h, 48h and 72h post-infection. (C) RT-qPCR of *iNos2, Gbp1*, and *Gbp5* transcript levels in Mtb infected WT and *Trim14* KO macrophages. (D) Proposed model of TRIM14’s dual roles in regulating cytosolic DNA sensing. TRIM14/cGAS interaction is ired to inhibit proteosomal degradation of cGAS and TRIM14/TBK1 interaction is required to promote TBK1-dependent phosphorylation of STAT3 at Ser727 and activate transcription of negative regulators of the type I IFN response like *Socs3*.

Because another report demonstrated that loss of TRIM14 leads to hyperreplication of VSV, an enveloped RNA virus, we also infected TRIM14 KO and control RAW 264.7 cells with VSV and followed viral replication and ISG expression by RT-qPCR over a 12-hour time-course. Although uptake of virus, as inferred by viral genome measurements at the 1h time point was very similar between the two genotypes, we observed a dramatic hyperreplication of VSV in *Trim14* KO macrophages (Fig. S4B) concomitant with significantly higher *Ifnb* and ISG expression (Figure S4D). Although hyperinduction of ISGs is seemingly at odds with hyperreplication of VSV, we do consistently observe low basal levels of *Ifnb* and ISGs in resting macrophages lacking TRIM14 (Fig. S4C) (likely due to cGAS instability (27)), which may give the virus enough time to “take off” before TRIM14-dependent resolution of type I IFN responses kicks in.

## DISCUSSION

To prevent chronic inflammation and damage to host cells and tissues, potent innate immune responses like type I IFN induction require tight temporal control. Here, we demonstrate a previously unappreciated role for TRIM14 in resolving *Ifnb* and ISG expression following a variety of cytosolic innate immune stimuli. By providing evidence that TRIM14 can directly interact with both cGAS and TBK1, our work uncovers a complex mechanism through which TRIM14 can both up and downregulate type I IFN responses in macrophages. Notably, we report that loss of TRIM14 has significant consequences on cell autonomous control of both bacterial and viral replication, with dramatic restriction of *M. tuberculosis* replication and uncontrolled replication of VSV observed in *Trim14* KO macrophages (Fig. 7A-B and S4). These results reveal a crucial role for TRIM14 in regulating macrophage innate immunity and point to TRIMs as potential targets for host-directed therapies designed to enhance a macrophage’s antimicrobial repertoire.

Our data support a model whereby TRIM14 acts as a scaffold between TBK1 and STAT3, promoting TBK1-dependent phosphorylation of STAT3 at Ser727 and transcriptional activation of negative regulators of JAK/STAT signaling like SOCS3 (Figure 7D). There is mounting evidence that a complex network of post-translational modifications regulates STAT3’s ability to dimerize, translocate to the nucleus, and/or bind DNA (41). In addition to inhibitory and activating STAT3 phosphorylation at Ser754 and Ser727 (42), respectively, several other modifications are known to control STAT3 activity, including acetylation at lysine 685 and phosphorylation of tyrosine 705, both of which increase the protein’s ability to bind DNA and translocate to the nucleus (56, 57). We propose that in the context of DNA sensing, TBK1-dependent phosphorylation of STAT3 acts as a control point for ramping up or down the STAT3 transcriptional regulon and the presence of TRIM14 can tip this balance. It is curious that these two modifications (Ser727 and Ser754) have dramatic opposing effects on STAT3 activity, as both residues reside in the transactivation domain in close proximity. Structural studies will be needed to shed light on how modulation of TBK1/STAT3 interactions by TRIM14 promote Ser727 phosphorylation over Ser754 phosphorylation. It is possible that the presence of TRIM14 makes one site more accessible either directly through interactions with STAT3 or by modulating interactions with other binding partners that influence availability of one serine over the other.

The apparent reliance of *Socs3* on STAT3 for its activation in our RAW 264.7 cells is also notable. In addition to being expressed by STAT3, depending on the cell type and context, *Socs3* can be transcribed by STAT1 and its promoter also contains AP-1, Sp3, and NFκB binding elements (58–60). The fact that these remaining transcription factors do not compensate for loss of STAT3 nuclear translocation in Trim14 KO macrophages (Fig. 5C) hints at the potential for crosstalk between STAT3 and other transcription factors, consistent with previous reports (61–63) and the extent to which the entire STAT3-transcriptional regulon is impacted by loss of TRIM14 remains unclear. Furthermore, following STAT3 expression of SOCS3, SOCS3 can actually downregulate STAT3 via a negative feedback loop (64, 65); future experiments will need to determine precisely how this loop is broken in *Trim14* KO macrophages. As STAT3 and SOCS3 are hugely important not only for controlling inflammatory responses during infection but also for regulating embryogenesis, cancer metastasis, and apoptosis, there is a critical need to understanding how TRIM14 can regulate their activation (66, 67).

Another recent study also shows a requirement for TRIM14 in VSV replication but reported that *Trim14* KO macrophages had lower ISG expression compared to wild-type (27). The authors ascribed these phenotypes to TRIM14’s role in promoting cGAS stabilization and provide evidence that loss of TRIM14 allows for cGAS degradation via the E3 ligase USP14 that targets cGAS to p62-dependent selective autophagy. We also observed lower *Ifnb* in response to *M. tuberculosis* and ISD transfection for our earliest measurements (4 hours for *M. tuberculosis* infection; 2, 4, and 6 hours for ISD transfection) (Fig. 3B, F, and G), but the phenotype of *Trim14* KO macrophages dramatically shifts to hyperinduction at later time points. It is not entirely clear what accounts for the discrepancies in our data, although notably, our analysis focuses almost exclusively on early time points following viral infection or innate immune activation (1-12 hours) as opposed to the 12-24 hours Chen *et al.* focused on, during which cell death resulting from high viral titers may complicate measurement and interpretation of transcript abundance. Taking the conclusions from both studies into account, it seems likely that TRIM14 plays a dual role in type I IFN regulation whereby it interacts with cGAS to promote type I IFN expression and with TBK1/STAT3 to dampen it (Fig. 7D). Future work will need to investigate the precise spatiotemporal distribution of cGAS/TRIM14-,TBK1/TRIM14-, and STAT3/TRIM14-containing complexes over the course of type I IFN induction and resolution. It will also be important to investigate how and when TRIM14 itself is post-translationally modified. Recent work from Jia *et al.*, provides evidence for RNF125-mediated polyubiquitination and proteosomal degradation of a mitochondrially-associated population of TRIM14 during viral infection (68). This post-translational modification and others could be critical for controlling whether TRIM14 influences type I IFN responses at the level of cGAS or TBK1/STAT3 or both.

Our finding that *Trim14* KO macrophages are better at controlling *M. tuberculosis* replication is quite remarkable. As *M. tuberculosis* replicates very slowly (~24-hour doubling time), we propose that unlike VSV, whose replication can be influenced by low resting ISGs in *Trim14* KO cells, *M. tuberculosis* replication is restricted by hyperinduction of ISGs that dominate after 4 hours of infection. It is unlikely that TRIM14’s contribution to cGAS stability accounts for *M. tuberculosis* restriction in *Trim14* KO macrophages, as our previous work showed that knocking out cGAS actually renders macrophages more permissive to *M. tuberculosis* infection, likely through loss of selective autophagy downstream of cytosolic DNA sensing (3). Consistent with these data and our model, another group has reported that siRNA knockdown of STAT3 in human macrophages enhances nitric oxide synthesis and restricts *M. tuberculosis* replication (69), although it is possible that TRIM14 contributes to *M. tuberculosis* restriction through more direct mechanisms as well. Curiously, in the context of *Listeria monocytogenes* infection of STAT1-deficient fibroblasts, overexpression of TRIM14 was protective, suggesting TRIM14 may have ISG-independent antibacterial functions (70). Future experiments designed to investigate what proteins TRIM14 interacts with on the *M. tuberculosis* phagosome and how loss of TRIM14 impacts maturation of the autophagosome will provide important insights into how TRIM14 controls *M. tuberculosis* replication and shed light on how we may be able to manipulate TRIM14 as a tuberculosis therapeutic.

## ACKNOWLEDGMENTS

We would like to acknowledge Monica Britton at the University of California, Davis DNA Technologies & Expression Analysis Core Library for her help with initial analysis of our RNA-seq data. We would also like to acknowledge the members of the Patrick and Watson labs for help with editing the manuscript, their valuable discussions, and critical feedback.

---

**Figure S1.**
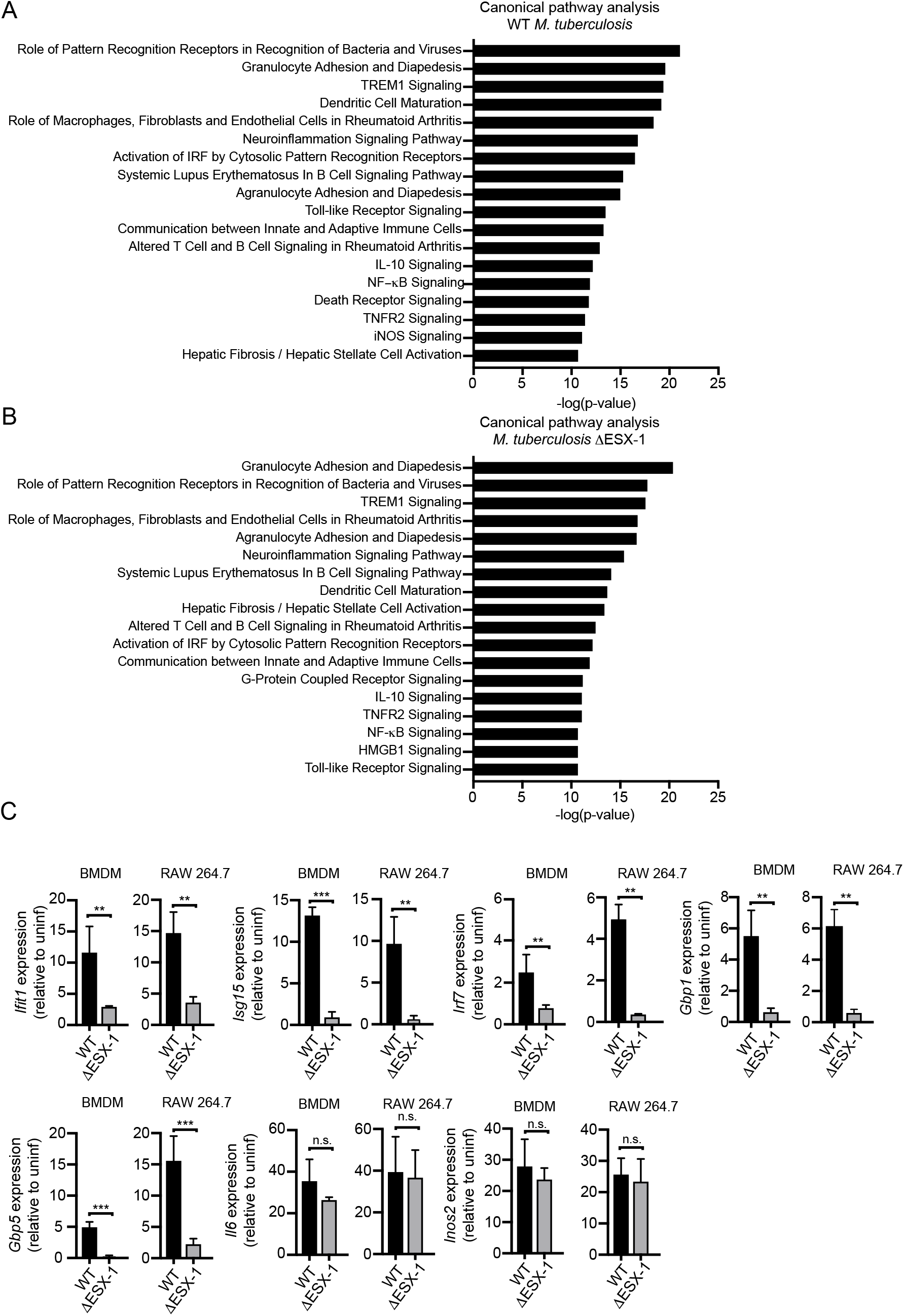
(A) IPA software analysis of cellular pathways enriched for differentially expressed genes in BMDMs infected with WT *M. tuberculosis*. (B) IPA software analysis of cellular pathways enriched for differentially expressed genes in BMDMs infected with ΔESX-1 *M. tuberculosis*. (C) RT-qPCR of transcripts in BMDMs & RAW 264.7 cells infected with WT *M. tuberculosis.*

**Figure S2.**
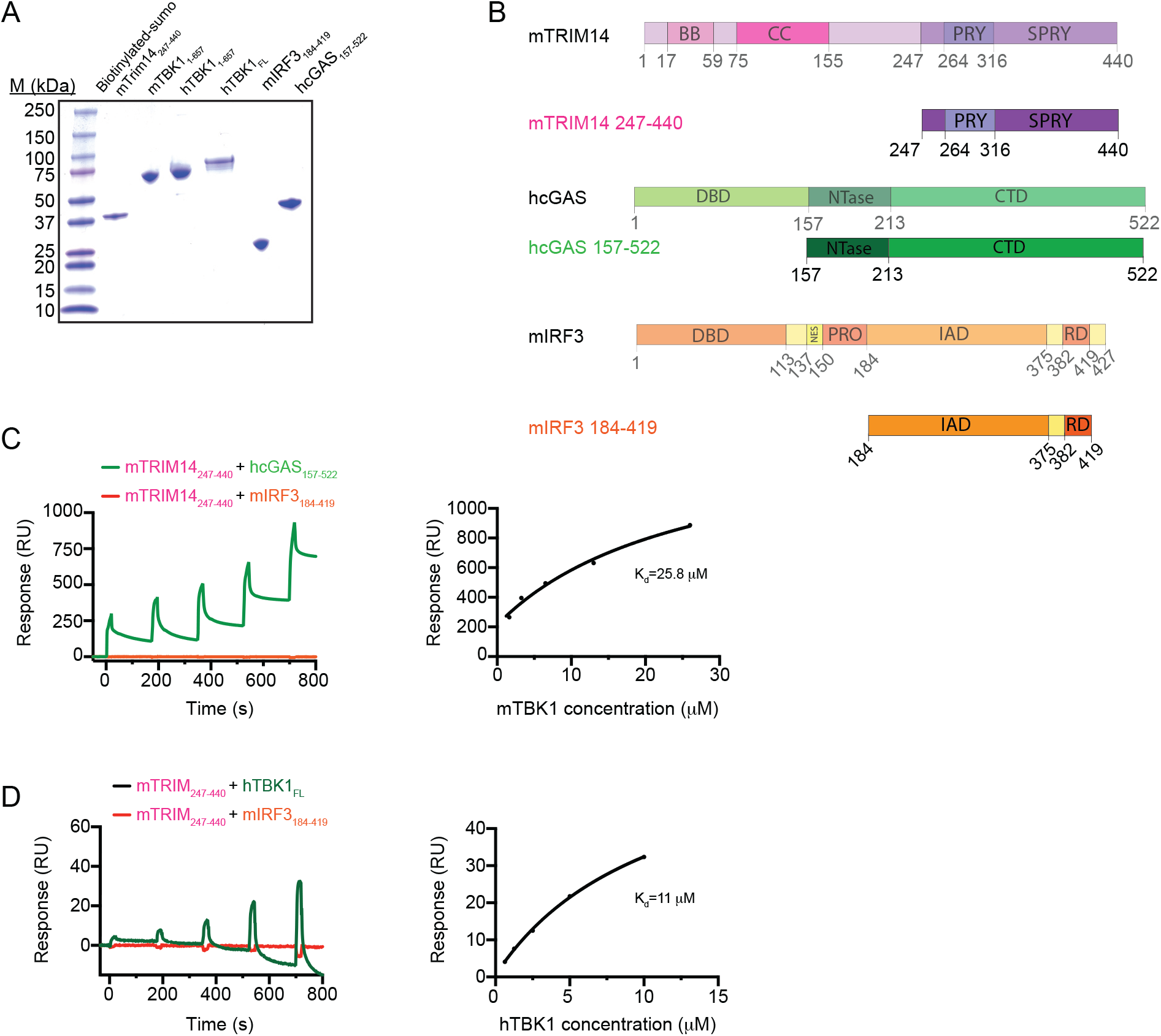
(A) Protein expression of mTRIM14, mTBK1, hTBK1, mIRF3, and hcGAS. (B) Diagram of mTRIM14, mIRF3, and hcGAS gene domains and truncations used in SPR studies. (C) Equilibrium binding study of mTRIM14 and hcGAS by surface plasmon resonance (SPR). mIRF3 was used as a negative control. Dissociation constant (K_d_= 25.8 μM) was derived by fitting of the equilibrium binding data to a one-site binding model. Equilibrium binding study of mTRIM14 and hTBK1 by surface plasmon resonance (SPR). mIRF3 was used as a negative control. Dissociation constant (K_d_= 11 μM) was derived by fitting of the equilibrium binding data to a one-site binding model.

**Figure S3.**
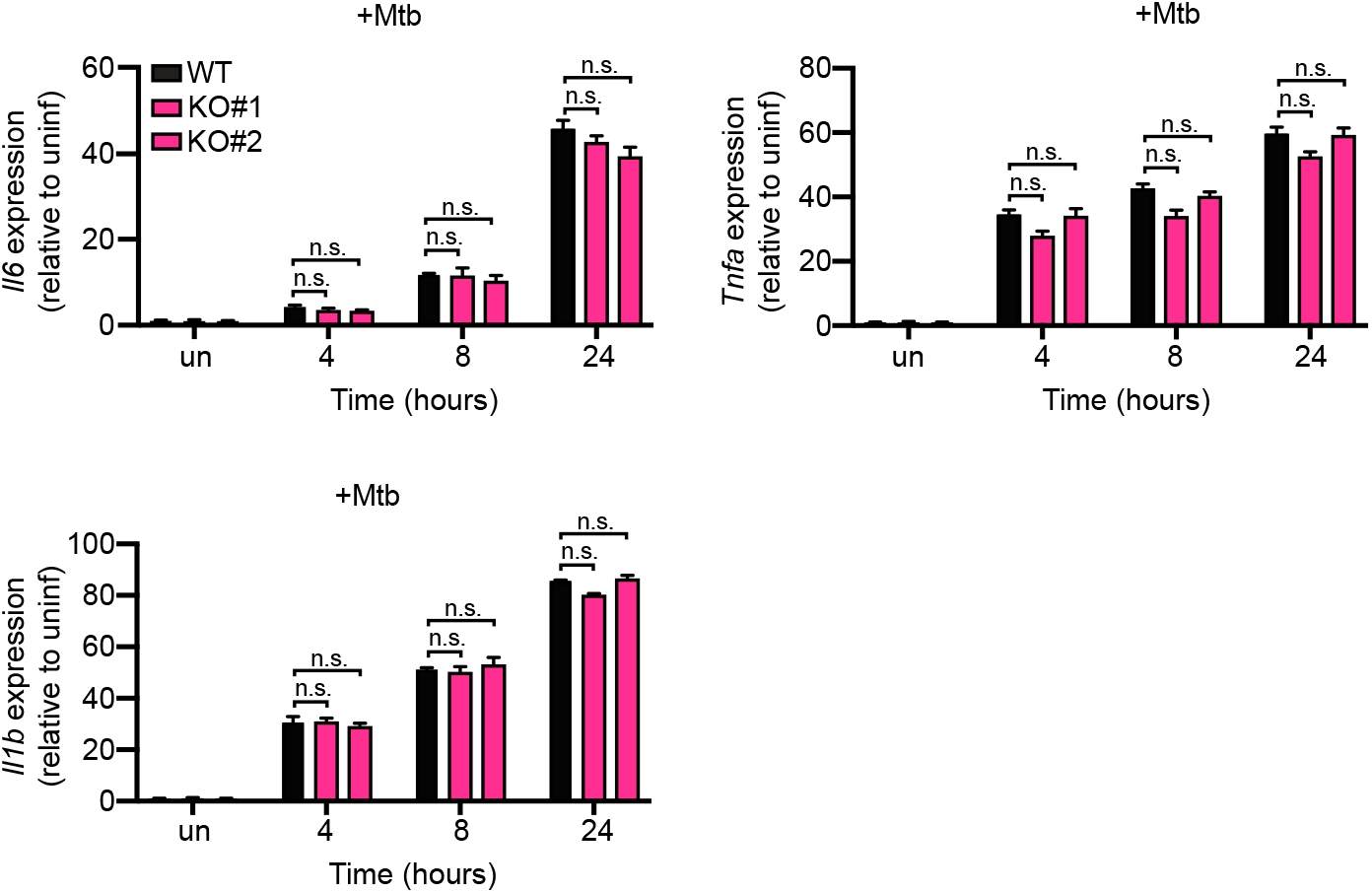
(A) RT-qPCR of *Il6, Tnfa,* and *Il1b* transcripts in WT and *Trim14* KO RAW 264.7 macrophages infected with *M. tuberculosis* at specified times after infection.

**Figure S4.**
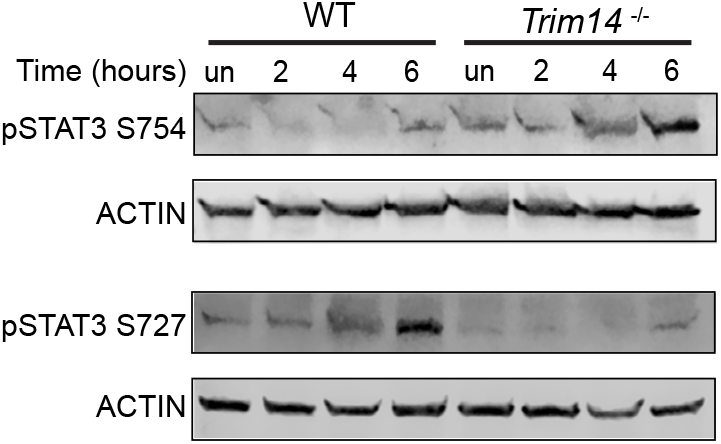
Immunoblot of phospho-STAT3 Ser754, phospho-STAT3 Ser727, and phospho-STAT1 Y701 in WT and *Trim14* KO RAW 264.7 macrophages at 1, 2, 4, 6, 8h post recombinant IFNb treatment. ACTIN is shown as a loading control.

**Figure S5.**
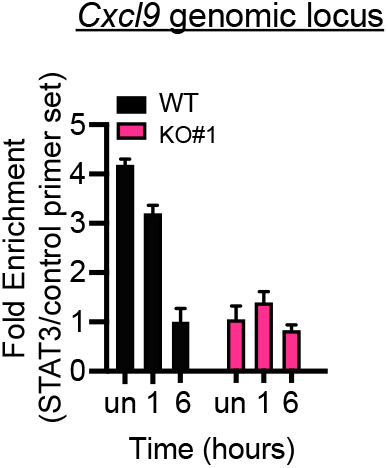
ChIP-qPCR of 3xFLAG-STAT3 associated genomic DNA from the CXCL9 locus in WT and *Trim14* KO RAW 264.7 macrophages transfected with 1 μg ISD.

**Figure S6.**
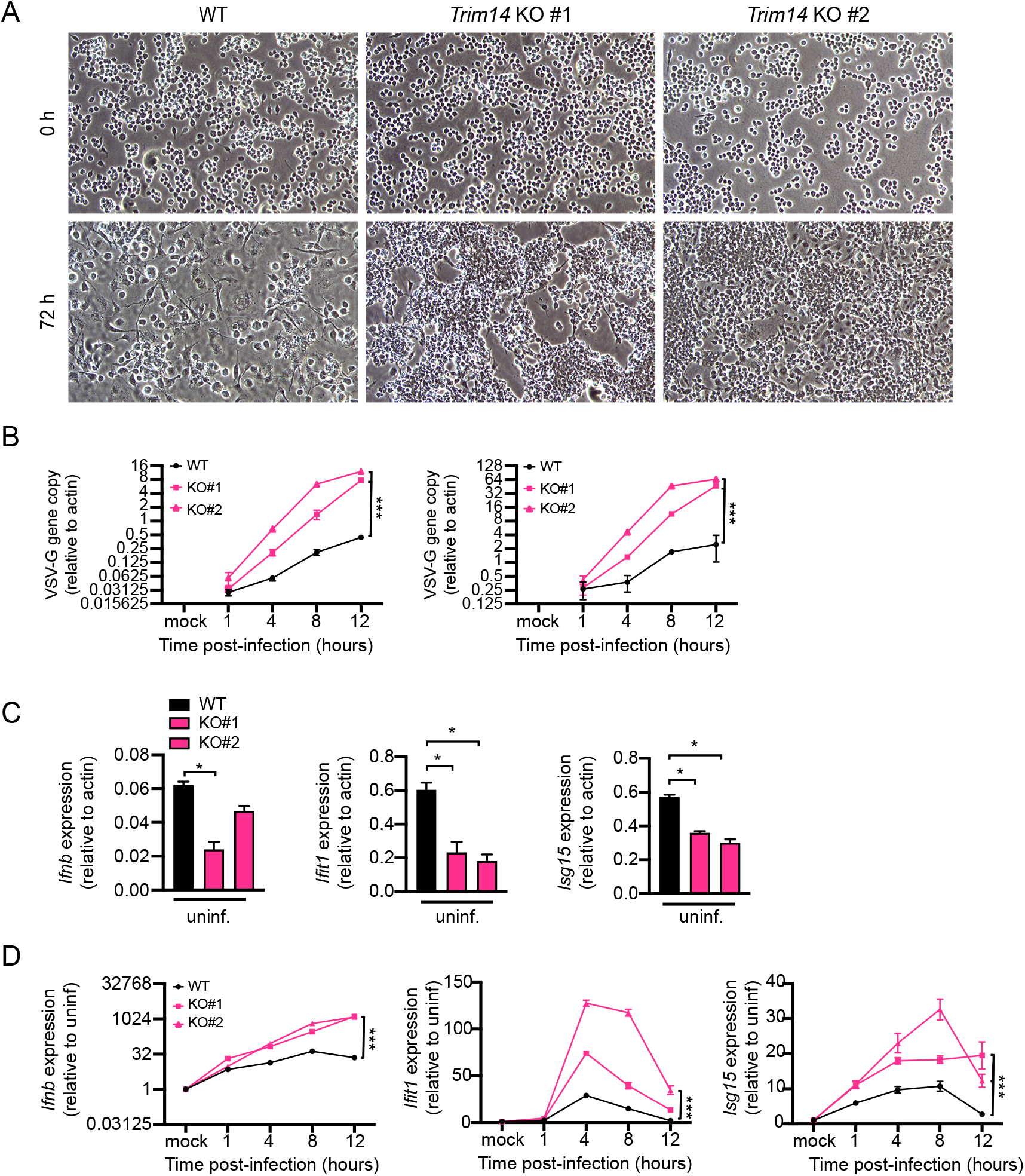
(A) Phase contrast images showing monolayers of WT and *Trim14* KO RAW 264.7 macrophages infected with Mtb *luxBCADE*. (B) Viral replication in WT and *Trim14* KO RAW 264.7 macrophages infected with VSV (MOI=1.0 or Mock infected) at 1h, 4h, 8h and 12h post-infection. (C) RT-qPCR of transcript levels in WT and Trim14 KO RAW 264.7 macrophages (D) RT-qPCR of transcript levels in WT and Trim14 KO RAW 264.7 macrophages infected with VSV (MOI=1.0 or Mock infected) at 1h, 4h, 8h and 12h post-infection.

